# Neuropilin-1 expression by solid tumors impairs CAR T cell function

**DOI:** 10.1101/2025.09.16.676545

**Authors:** Louise Condon, Marie Bouillié, Jaime Fuentealba, Michael Saitakis, Olivier Hermine

## Abstract

Neuropilin-1 (NRP-1) is a versatile transmembrane protein expressed in numerous cell types and tissues, both in health and in disease. In particular, it is expressed by activated T cells, where it has been shown to play an inhibitory role, dampening their response to tumor cells. Chimeric antigen receptor (CAR) T cells have had considerable success in treating cancers such as B cell leukemias, but face several obstacles in the treatment of solid tumors.

We hypothesized that NRP-1 may contribute to dampening CAR T cell responses to cancer. While NRP-1 blockade either through neutralizing antibodies or through CRISPR/Cas9-mediated deletion did not improve CAR T cell cytotoxic activity in an *in vitro* solid tumor model, NRP-1 was found to have a protective effect against CAR T cells when expressed by the tumor cells, in line with previous literature implicating NRP-1 as a pro-tumor factor involved in their growth and invasiveness.

We found that NRP-1 was transferred from target cells to CAR T cells via trogocytosis, i.e. the “nibbling” and subsequent display of membrane proteins from one cell by another. CD19, the target antigen in this model, was also transferred to CAR T cells, raising interesting questions about trogocytosis as a marker for effective tumor cell killing through direct interaction. On the contrary, trogocytosis may impede effective CAR T function by promoting fratricide due to their display of target antigen.

While no conclusive mechanisms for NRP-1 activity were found, several avenues of research have been opened up to further our understanding of the multifaceted roles of NRP-1 in CAR T cell activity and interactions with their targets.

## Introduction

Neuropilin-1 (NRP-1) is a 130 kD transmembrane glycoprotein that is expressed in multiple cell types and involved in multifaceted processes, including angiogenesis, cell adhesion and migration, cancer and immunity (Soker *et al*., 1998; Romeo *et al*., 2002; Ellis, 2006; Chaudhary *et al*., 2014; Roy et al., 2017). It interacts with many ligands from distinct molecular families, including class 3 semaphorins (He and Tessier-Lavigne, 1997) and growth factors such as vascular endothelial growth factor (VEGF) (Soker *et al*., 1998) and transforming growth factor β (TGFβ) (Glinka and Prud’homme, 2008).

NRP-1 is expressed at the plasma membrane in dendritic cells as well as in T lymphocytes and is upregulated in activated T cells and regulatory T cells (Tregs) (Bruder et al., 2004; Sarris et al., 2008; Rossignol et al., 2022). It polarizes at the immune synapses that are formed between CD8_+_ T cells and dendritic cells (DCs) or tumor cells, and has been identified as an inhibitory checkpoint molecule in these effector cells. NRP-1 cooperates with programmed death 1 (PD-1) at the immune synapse to dampen T cell responses to tumor cells, *in vitro* and in mouse models, inhibiting effective anti-tumor immunity (Leclerc *et al*., 2019; Rossignol *et al*., 2022). In particular, binding of NRP-1 to one of its main ligands, semaphorin 3A (Sema3A), causes impaired synapse polarization and Zeta-chain-associated protein kinase 70 (ZAP-70) phosphorylation, an essential irst step in the T cell activation cascade (Rossignol *et al*., 2022). NRP-1 has also been shown to inhibit T cell memory differentiation, leading to poor long-term anti-tumor immune responses (C. Liu *et al*., 2020). NRP-1 blockade with monoclonal antibodies, as well as NRP-1 ablation by gene editing, both resulted in an invigorated anti-tumoral T cell response when combined with anti-PD-1 therapies (Leclerc *et al*., 2019; Rossignol *et al*., 2022).

While a few studies have now established the inhibitory role of NRP-1 in conventional T cells, there is as yet no indication of the role it might play in chimeric antigen receptor (CAR) T cells. CAR T cells are genetically engineered T cells expressing a specific receptor for one or several tumor antigens, allowing them to recognize and attack tumor cells that might otherwise evade the immune system. CAR T cell therapy has shown considerable success in treating hematological cancers such as B-cell acute lymphoblastic leukemia (B-ALL) and diffuse large B-cell lymphoma (DLBCL), by targeting a specific B cell antigen, CD19 (Sermer et al. 2020; Pasvolsky et al. 2023). However, CAR T cell therapy must still surmount several challenges to fulfill its vast potential. CAR T cells have been much less successful in treating solid tumors, as the tumor sites are typically harder to penetrate due to a hostile physical, chemical, and cellular environment. Dense, chaotic extracellular matrix; inhibitory cytokines and chemokines; and exhaustion-inducing ligands expressed by immunomodulatory cells in the tumor niche are all obstacles to CAR T cell infiltration and function within the tumor microenvironment (G. Liu *et al*., 2021). Additionally, the search for new antigenic targets is ongoing, requiring both high expression by tumor cells and low expression by surrounding healthy tissues, to avoid off-target toxicity. Antigenic escape is common in many cancers, usually through adaptive downregulation of target antigen expression, or through natural selection of the least immunogenic clones in a heterogeneous tumor (Majzner and Mackall, 2018). Another, less well-known mechanism for loss of tumor antigens is antigen capture by CAR T cells *via* a process called trogocytosis.

Trogocytosis, or cell “nibbling”, is a process resembling partial endocytosis or phagocytosis and employing some of the same molecular machinery (Hudrisier *et al*., 2001). In vertebrates, this behavior has been observed in natural killer cells, monocytes, and other cells of the immune system as a mechanism of pathogen killing (Bettadapur *et al*., 2020). It can occur upon immunological synapse formation when plasma membrane fragments and proteins are pulled from the target cell by the effector cell and displayed on the effector cell membrane. Trogocytosis plays several important and sometimes contradictory roles in the immune response to pathogens and to cancer cells: it can act as a mechanism to spread foreign and tumor antigens for rapid and efficient antigen presentation; conversely, it can promote tumor escape by stripping antigens from tumor cells, reducing antigen density and thus their immunogenicity; it can also cause fratricide among CAR T cells that recognize their target antigen on each other (Hamieh *et al*., 2019; Schoutrop *et al*., 2022). NRP-1 has been shown to transfer between cells *via* trogocytosis in the similar context of the dendritic cell (DC)-T cell synapse (Bourbié-Vaudaine *et al*., 2006).

While CAR T cells were designed to recreate canonical T cell activation mechanisms, using preexisting biological machinery downstream of the CAR as well as within the CAR structure, mounting evidence suggests that the biology of CAR T cells may diverge more fundamentally from conventional T cell biology than previously thought. The intensity and duration of CAR T cell activation, for example, appear to differ significantly from TCR-mediated T cell activation, due to differences in affinity and costimulatory domains, among other factors (Teppert *et al*., 2022).

In this study, we aim to elucidate the possible roles of NRP-1 in these uniquely specialized cells, and whether it fulfills similar functions as those described in conventional T cells. We also study the effects of tumor NRP-1 expression on CAR T cell function and explore some potential mechanisms for this versatile protein’s impact on CAR T cell-mediated anti-tumor immunity.

## Results

### Neuropilin-1 is expressed in stimulated T cells

We irst aimed to verify whether NRP-1 was indeed expressed in classically activated T cells *in vitro*. To do so, healthy donor-derived peripheral blood mononuclear cells (PBMCs) were sorted negatively to isolate CD3^+^ cells. These cells were then stimulated using magnetic beads carrying anti-CD3 and anti-CD28 antibodies; fresh beads were added on a weekly basis and were removed magnetically after 3 to 4 days of contact with the cells. The activated T cells were periodically stained with fluorescent antibodies and analyzed by flow cytometry.

While NRP-1 expression was not detectable prior to stimulation, the protein was detected at low levels after the first stimulation and was increasingly highly expressed after each restimulation (**Figure 1a** and **1b**). Indeed, up to 70% of total T cells expressed NRP-1 at peak, a proportion considerably higher than previously reported. Previous research from Leclerc *et al*. (2019) and Rossignol *et al*. (2022) found that NRP-1 was expressed almost exclusively by activated PD-1^+^ cells. Unexpectedly, we identified a cell population expressing NRP-1 but not PD-1, and this NRP-1^+^ PD-1^-^ population increased after subsequent restimulations and rest periods (**Figure 1c**), indicating that NRP-1 was not always co-expressed with PD-1, as previously suggested.

**Figure 1.**
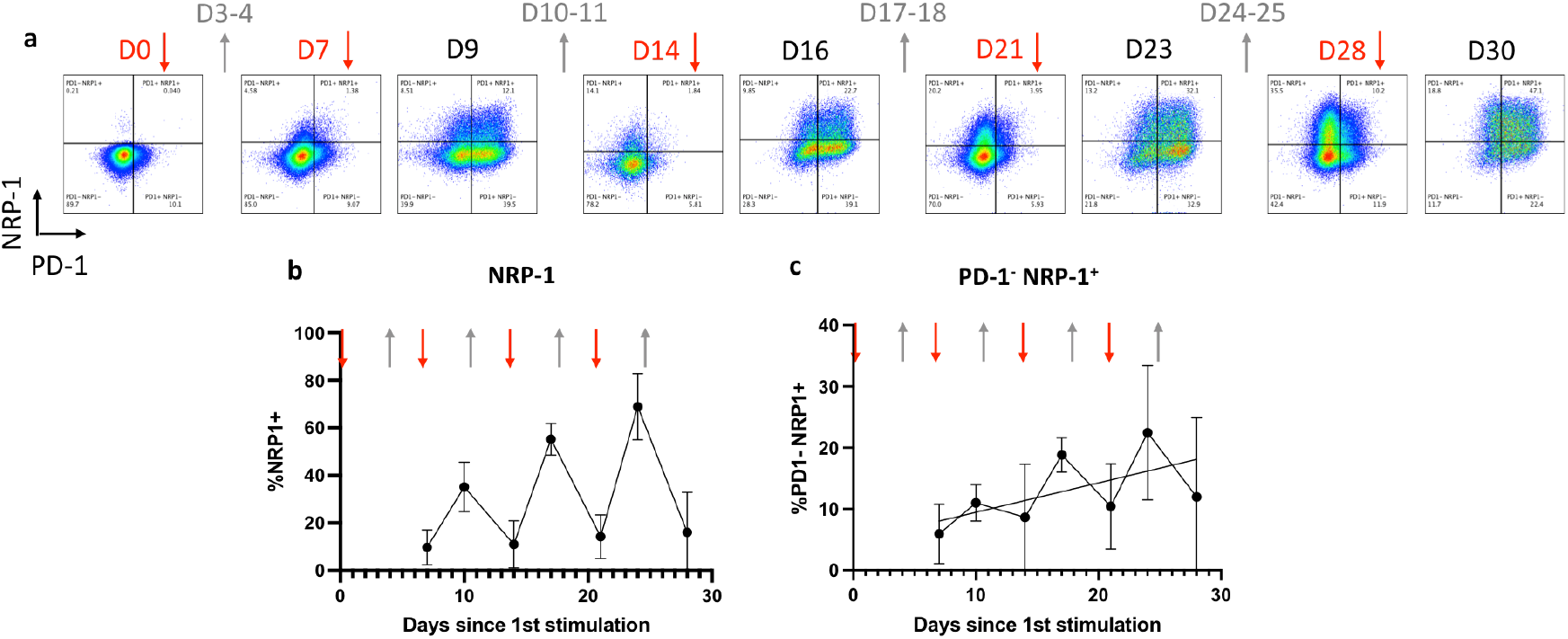
**a.** Representative kinetics of NRP-1 and PD-1 expression. Red arrows represent addition of fresh anti-CD3/anti-CD28 beads, grey arrows represent removal of the beads from the medium. **b**. Average percentage of cells expressing NRP-1 over time with repeated stimulations, from 5 independent experiments. **c**. Average percentage of PD-1-NRP-1^+^ cells over time with repeated stimulations, from 5 independent experiments.

These expression kinetics were identical in CAR T cells stimulated in the same way, by targeting the TCR with anti-CD3/anti-CD28 magnetic beads (data not shown). Similar levels of NRP-1 expression occurred in transduced and non-transduced T cells, both before and after restimulation, indicating that NRP-1 expression was not induced by the CAR construct itself.

### Anti-NRP1 antibodies do not alter CAR T cell killing of NRP1^+^ target cells

Leclerc *et al*. (2019) showed that NRP-1 blockade using a monoclonal antibody could enhance the CD8^+^ T cell response to tumor cells in mice, especially in combination with anti-PD1 antibodies.

We therefore sought to determine whether NRP-1 blockade could likewise enhance the CAR T cell response to human tumor cell lines *in vitro*. Specifically, we explored CAR T efficacy in a solid tumor model by culturing target cell lines derived from solid tumors in spheroid form. This method more closely mimics the solid, three-dimensional tumors found *in vivo* than two-dimensional culture in flat-bottom wells.

Healthy primary CD3^+^ cells were stimulated with anti-CD3/anti-CD28 magnetic beads, then transduced with a lentivirus carrying a commercial CD19-BBz CAR construct. CAR expression was ascertained using flow cytometry and was shown to be stable over several weeks in 60 to 75% of total cells, depending on the blood donor (**Figure 2a**).

**Figure 2.**
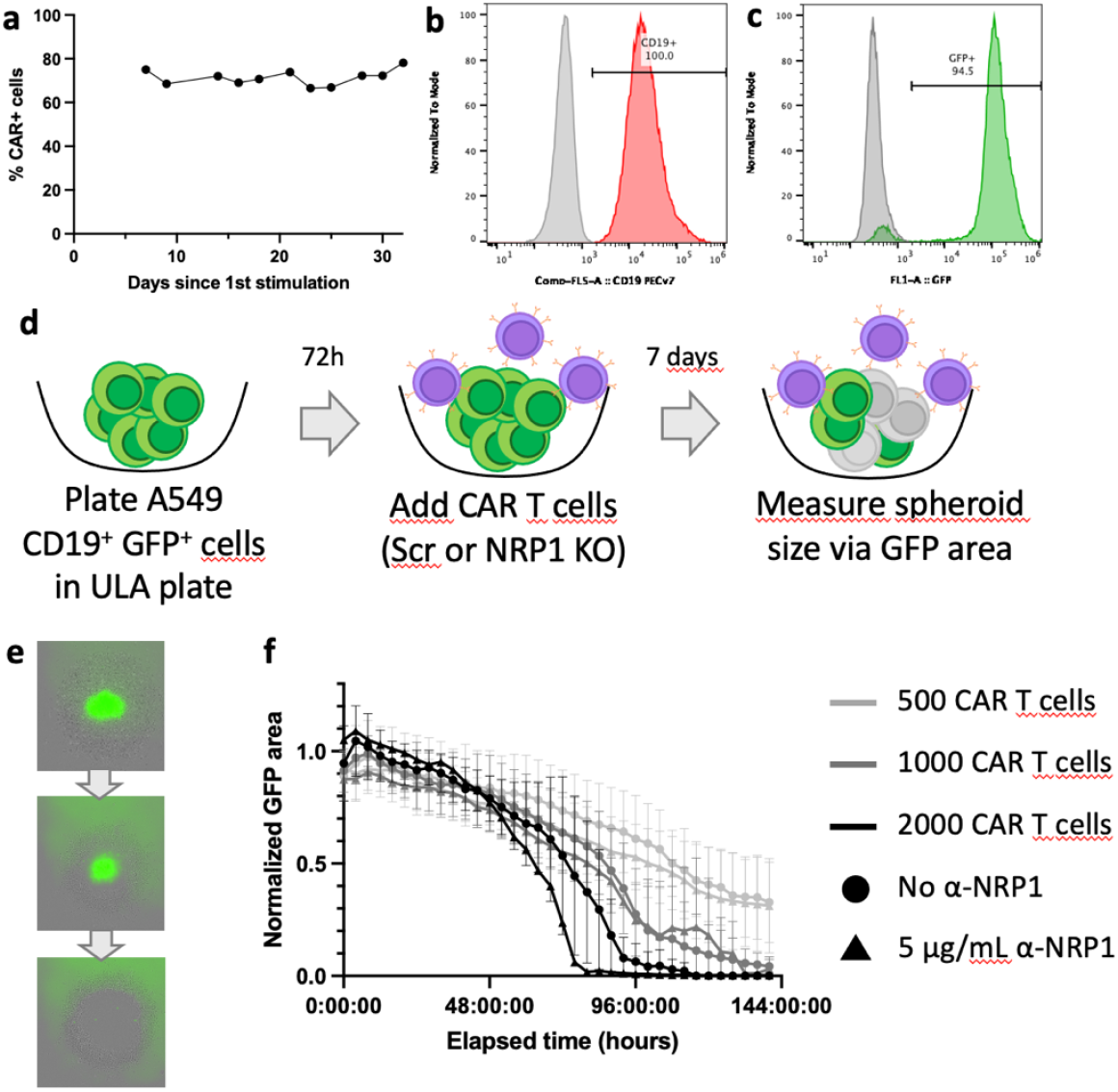
**a.** Representative CAR expression over time, with repeated stimulations. **b**. Representative A549-CD19 surface expression of CD19. **c**. Representative A549-CD19 expression of GFP. **d**. Schematic spheroid assay experimental setup. **e**. Representative evolution of an A549-CD19 spheroid co-cultured with a high dose of CAR T cells. **f**. Change in GFP area in real-time cell monitoring, according to CAR T cell dose and addition of NRP-1-neutralizing antibodies. Data from 3 independent experiments, normalized to spheroid size in the absence of CAR T cells in each experiment. Bars indicate standard errors.

Next, human lung carcinoma cell line A549 was transduced stably with CD19, thus presenting an appropriate target for the CAR T cells. They were also transduced separately with GFP. The expression of CD19 and of GFP were confirmed in over 95% of cells by flow cytometry (**Figure 2b** and **2c**).

To evaluate CAR T cell efficacy against these tumor cells, we irst seeded 1000 A549-CD19 cells in 96-well ultra-low adhesion (ULA) plates, allowing them to grow undisturbed for 3 days, during which time they spontaneously formed three-dimensional spheroids.

Varying doses of CAR T cells were then seeded onto these spheroids. The resulting co-cultures were monitored in real time, imaging each well at 4-hour intervals using an IncuCyte machine (**Figure 2d**). The A549-CD19-GFP spheroids appear green in these images; upon contact with CAR T cells, this green area representing the live target cells gradually shrunk over the period of a week. When co-cultured with a high dose of CAR T cells, the green area disappeared entirely within as little as three days (**Figure 2e**). Using cytometry to analyze the contents of these wells, we ascertained that GFP^+^ cells were absent, and that only effector cells remained alive. Thus, the GFP area observed in real-time cell monitoring was a good proxy for A549-CD19 target cell survival.

In all future experiments, unless otherwise specified, the size of the spheroids in co-culture with CAR T cells is normalized to the unimpeded growth of the same target cells in the absence of effector cells: relative spheroid size is expressed as normalized GFP area.

In order to determine whether NRP-1 blockade could impact CAR T cell efficacy against A549-CD19 spheroids, target cells were seeded three days prior (D-3), then effector cells were added at D0. 5 µg/mL anti-NRP1 were added to the test conditions, and control conditions had no anti-NRP1 antibody added.

Surprisingly, NRP-1 blockade had no effect on CAR T cell cytotoxicity in this model (**Figure 2f**). A trend towards enhanced CAR T cell efficacy was visible at the highest dose of effector cells (2000 CAR^+^ cells per spheroid), but this was not statistically significant.

While NRP-1 expression was measured in both target and effector cells before co-culture, it was not monitored kinetically over the course of the experiment. However, in subsequent experiments, we showed that NRP-1 expression increased in CAR T cells following contact with A549-CD19 target cells, and that NRP-1 expression in target cells was stable over the course of these co-cultures. This confirms that the anti-NRP-1 antibodies had an available antigenic target, but did not appear to have any effect on CAR T cell efficacy despite NRP-1 expression.

### NRP-1 knock-out in CAR T cells has no effect on their anti-tumor efficacy

A549 cells express NRP-1 constitutively and can thus be targeted by anti-NRP-1 antibodies. Our initial model for NRP-1 blockade did not account for this alternative target effect, leading us to opt for a more specific inhibition of NRP-1 expression in CAR T cells, by CRISPR-Cas9-mediated NRP-1 knock-out (KO).

To generate these NRP-1 KO CAR T cells, CD3^+^ cells were stimulated and transduced with CAR lentivirus as described before, then transfected with ribonucleoprotein complexes containing Cas9 protein and a mixture of three guide RNAs targeting the NRP-1 gene (**Figure 3a**). Alternatively, control CAR T cells were transfected with either Cas9 alone (Mock), or Cas9 alongside a scrambled gRNA not targeting any known human DNA sequence (Scr). Both control methods yielded similar levels of NRP-1 expression under identical conditions and similar results in subsequent experiments, and are thus used interchangeably.

**Figure 3.**
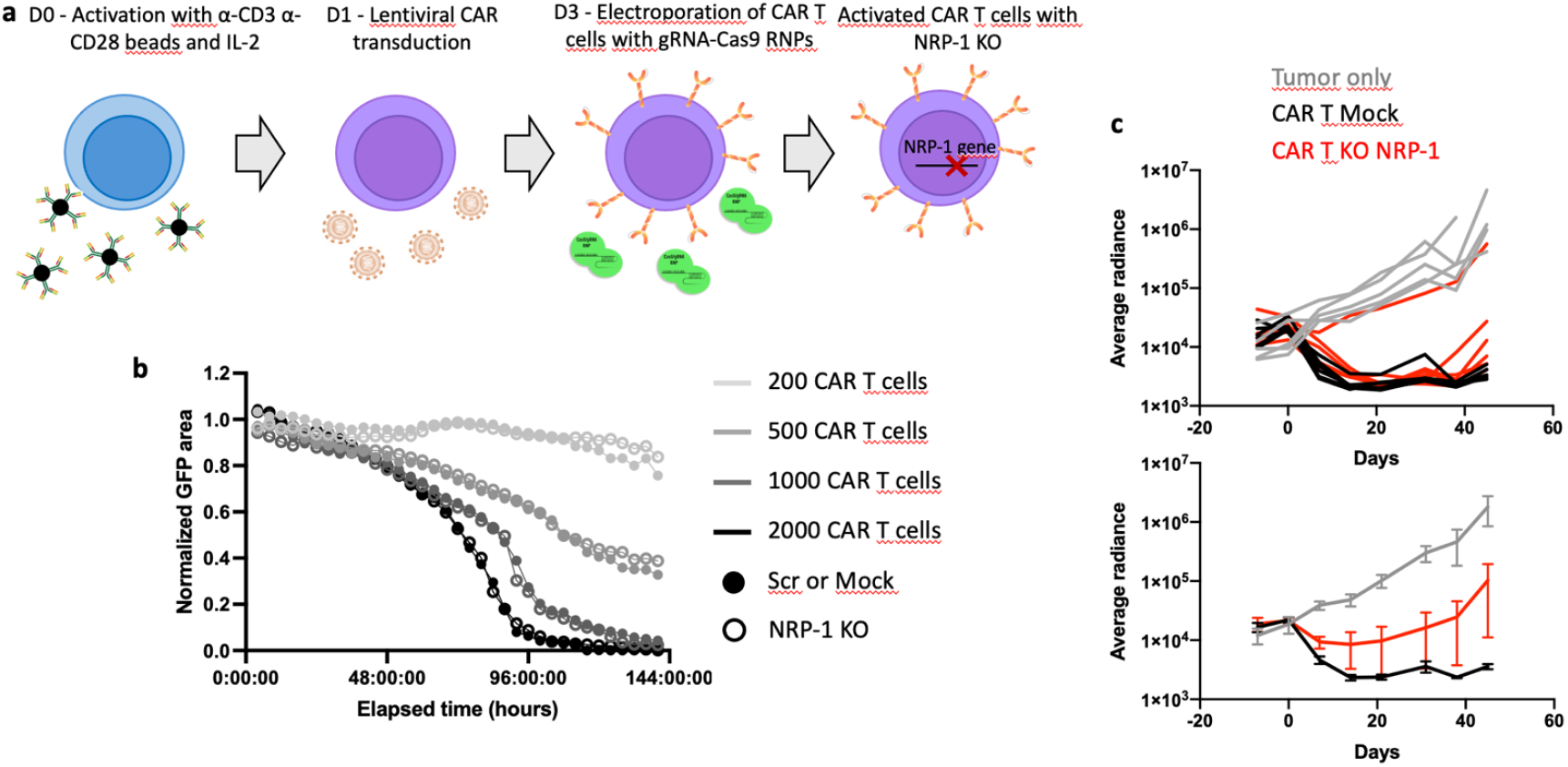
**a.** Experimental setup for NRP-1 KO CAR T cell production. **b**. Evolution of the spheroid GFP area in real-time cell monitoring, according to CAR T cell dose and NRP-1 KO in the CAR T cells. Data from 6 independent experiments, normalized to spheroid size in the absence of CAR T cells in each experiment. **c**. A549-CD19 tumor radiance in NSG mice treated with Mock or NRP-1 KO CAR T cells (individual mice in upper panel, averages and standard error bars in lower panel).

As before, A549-CD19 spheroids were seeded in ULA plates, followed three days later by the addition of different doses of either Scr or NRP-1 KO CAR T cells. Spheroid area decreased in a dose-dependent manner, disappearing entirely within four days at the highest CAR T cell dose. However, no differences in the rate of spheroid death were observed between the NRP-1 KO and Scr CAR T conditions, indicating that the lack of NRP-1 in effector cells did not improve their efficacy *in vitro* (**Figure 3b**).

We next asked whether these NRP-1 KO CAR T cells would be more efficient than Mock CAR T cells *in vivo*. Young adult NSG mice were injected intravenously with luciferase-expressing A549-CD19 cells, which were left to invade the lungs and form tumors for 21 days. Different doses of Mock or NRP-1 KO CAR T cells were then injected intravenously, and tumor size was evaluated on a weekly basis by administering luciferin intraperitoneally and imaging the mice for tumor luminosity. Almost all tumors were rapidly controlled by the CAR T cells, with no significant differences in their shrinkage rate (**Figure 3c**). However, relapse was observed in a few animals around 45 days after CAR T cell injection, and appeared more extensive in those that had received NRP-1 KO CAR T cells, in contrast to the continuously suppressed tumors in mice that received Mock CAR T cells. This indicates that *in vivo* also, NRP-1 KO does not improve CAR T cell efficacy, and may even result in an increased risk of tumor relapse.

### NRP-1 expression by tumor cells is protective against CAR T cells

Given that NRP-1 is expressed in both target cells and effector cells, we wondered whether A549-specific NRP-1 blockade could impact CAR T cell efficacy. Extensive research has shown that tumor NRP-1 expression is associated with poor prognosis in many cancer types (reviewed by Ellis, 2006). Several roles have been suggested to explain how NRP-1 may be beneficial to tumors, including its function in angiogenesis and in migration, which could increase tumors’ blood and oxygen supply as well as encourage metastasis. However, no published data have proven a direct role for NRP-1 in tumor resistance to CAR T cell therapy.

To answer this question, we generated a NRP-1 KO A549-CD19 cell line using CRISPR-Cas9 editing. This cell line did not significantly differ from the previously described A549-CD19 cell line in terms of growth rate, CD19 or GFP expression, or any other visible phenotypical parameters (**Figure 4a**). However, when cultured in spheroid form, a visual difference was observed in the density of the agglomerated cells (**Figure 4b**): A549-CD19 NRP-1^-^ cells formed looser spheroids than NRP-1^+^ cells, possibly facilitating the infiltration of cytotoxic cells in subsequent experiments. Interestingly, anti-NRP1 antibodies had no visible effect on spheroid density, possibly due to being added after the spheroids were formed.

**Figure 4.**
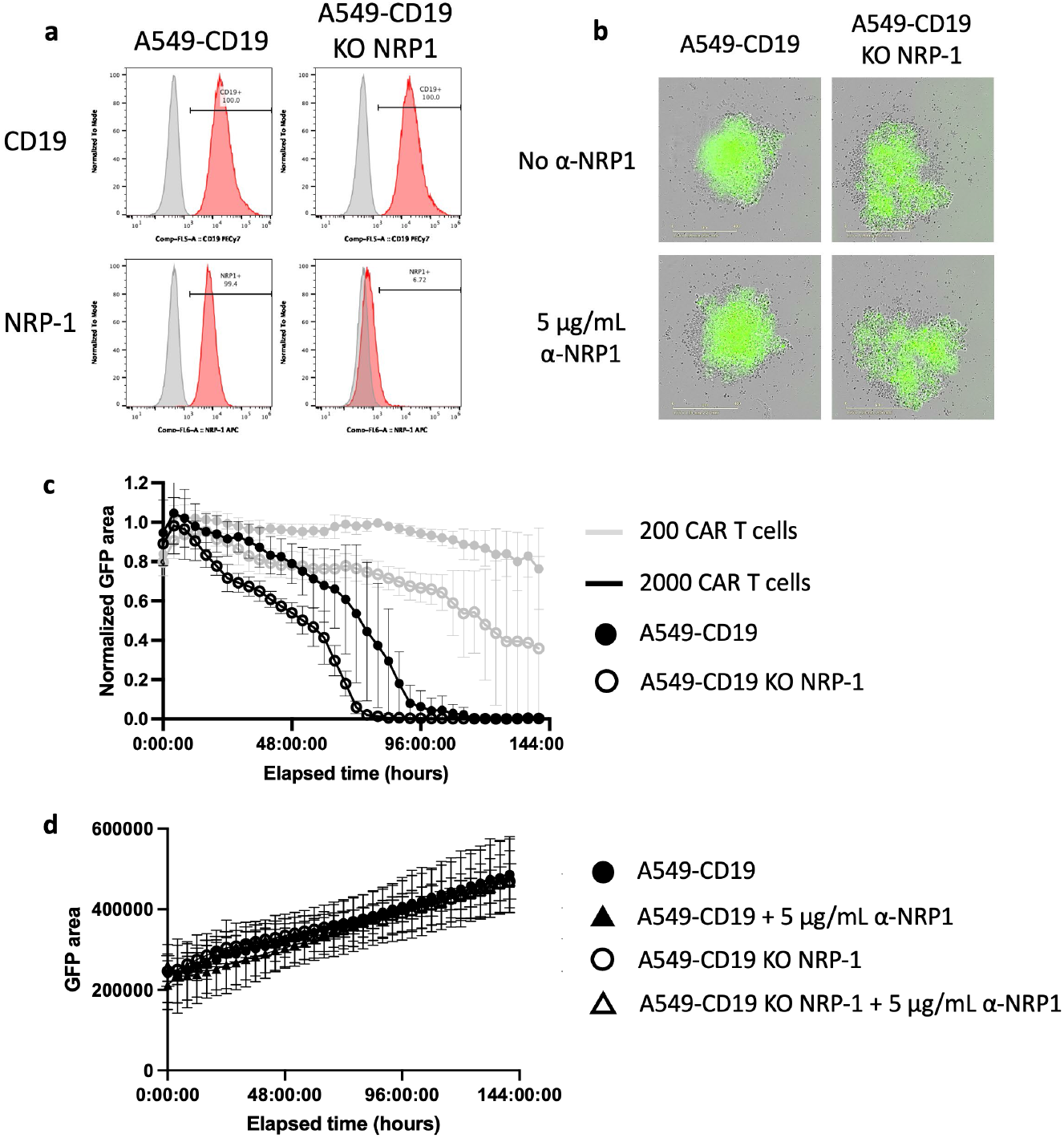
**a.** CD19 and NRP-1 expression in A549-CD19 and A549-CD19 KO NRP-1 cell lines. Grey: isotype control; red: CD19 or NRP-1 antibody. **b**. Representative A549-CD19 and A549-CD19 KO NRP-1 spheroids, both expressing GFP, after a week of growth. 5 µg/mL α-NRP1 was added to the medium at day 3 in the bottom panels only. Merged brightfield and green channel images. **c**. Evolution of the spheroid GFP area in real-time cell monitoring, according to CAR T cell dose and NRP-1 KO in the target cells. Data from 3 independent experiments, normalized to spheroid size in the absence of CAR T cells in each experiment. Bars indicate standard errors. **d**. Evolution of the spheroid GFP area in real-time cell monitoring, in the absence of CAR T cells, with two methods of NRP-1 blockade, genetic deletion and NRP-1-neutralizing antibodies. Data from 5 independent experiments, without normalization.

When co-cultured in spheroid form with CAR T cells, the NRP-1^-^ target cells were killed significantly more efficiently than the NRP-1^+^ cells, with the GFP area disappearing a full day earlier at the highest effector cell dose (**Figure 4c**).

This disparity was only observed in the presence of CAR T cells: spheroids growing alone and unimpeded increased in size at the same rate, with or without NRP-1 blockade (**Figure 4d**).

Thus, NRP-1 expression by cancer cells appears to have a protective effect against CAR T cells in this model. This may be linked to the tighter structure of NRP-1^+^ spheroids potentially impeding CAR T cell infiltration; or given the many roles of NRP-1 in cell-cell interactions, other mechanisms could be at play in reducing CAR T cell efficacy against NRP-1^+^ tumors.

When evaluating the effect of NRP-1 genetic deletion in CAR T cells, no change in killing ability was found (**Figure 3b**), whereas it was slightly enhanced using anti-NRP1 and a high dose of CAR T cells (**Figure 2f**). This raises the question of whether genetic deletion and antibody-mediated blockade of NRP-1 could have different consequences. To assess the specific effects of NRP-1 antibody blockade on CAR T cells requires studying their cytotoxicity in the presence and absence of anti-NRP1 antibodies, this time against A549-CD19 KO NRP-1 target cells, which are indifferent to the presence of anti-NRP1. However, NRP-1 antibody blockade in CAR T cells had no effect on their ability to lyse A549-CD19 NRP-1 KO target cells (data not shown). In this model, NRP-1 blockade using antibodies and genetic deletion respectively had indistinguishable effects on CAR T cell activity.

Thus, the precise effects of NRP-1 blocking antibodies on A549-CD19 target cells remain elusive and may involve complex mechanisms and interactions with CAR T cells.

We next attempted to elucidate the mechanisms for the protective function of NRP-1 in cancer cells, since understanding the causes for its deleterious effects against CAR T cell activity could yield insight on new strategies to combat them.

### NRP-1 is transferred from target cells to CAR T cells via trogocytosis

We noticed that many NRP-1 KO CAR T cells still displayed NRP-1 at the plasma membrane, detectable in flow cytometry, after co-culture with A549-CD19 target cells (**Figure 5**). This occurred despite almost total NRP-1 knock-out in the bulk T cell population, which consistently remained under 2% of total cells expressing NRP-1 when cultured alone and restimulated with anti-CD3/anti-CD28 beads. One possible explanation for this observation was that the very small NRP-1^+^ subpopulation was being heavily selected for during co-culture due to intense evolutionary pressures from the environment. However, when the effector and target cells were co-cultured for shorter time periods, NRP-1 was detected on the effector cells after only 24 hours, a timeframe too short for such a dramatic expansion of the rare NRP-1^+^ population. We concluded it was more likely that the NRP-1 molecules found on the NRP-1 KO CAR T cell membranes were coming from another source, namely the NRP-1^+^ target cells.

**Figure 5.**
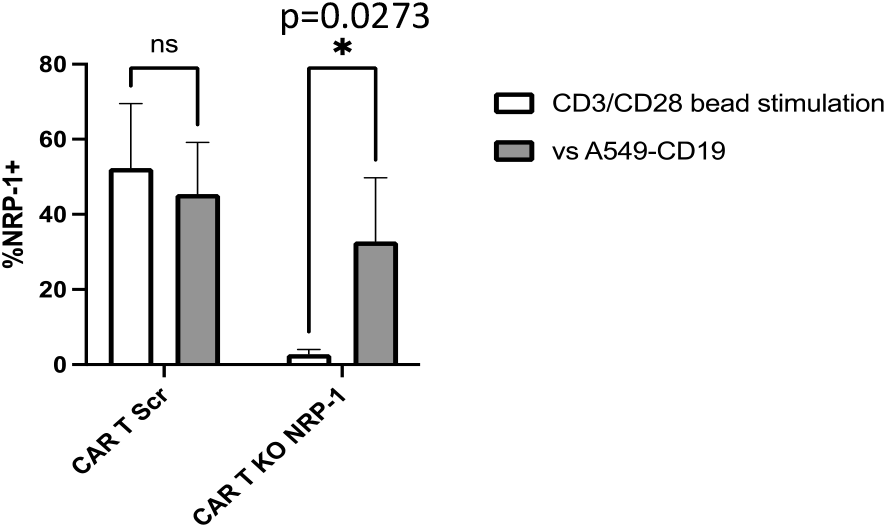
NRP-1 expression on NRP-1 KO CAR T cells after co-culture with A549-CD19 target cells or after *in vitro* stimulation with beads.

To positively verify whether NRP-1 was being transferred from the target cells, NRP-1 KO CAR T cells were co-cultured with A549-CD19 NRP1 KO target cells, eliminating this possible source of NRP-1 for the CAR T cells. In this condition, almost no NRP-1 was detectable on the CAR T cells by flow cytometry (**Figure 6a**, bottom panels, and **6b**). This indicates that NRP-1 is indeed transferred from the target to the effector cells during co-culture.

**Figure 6.**
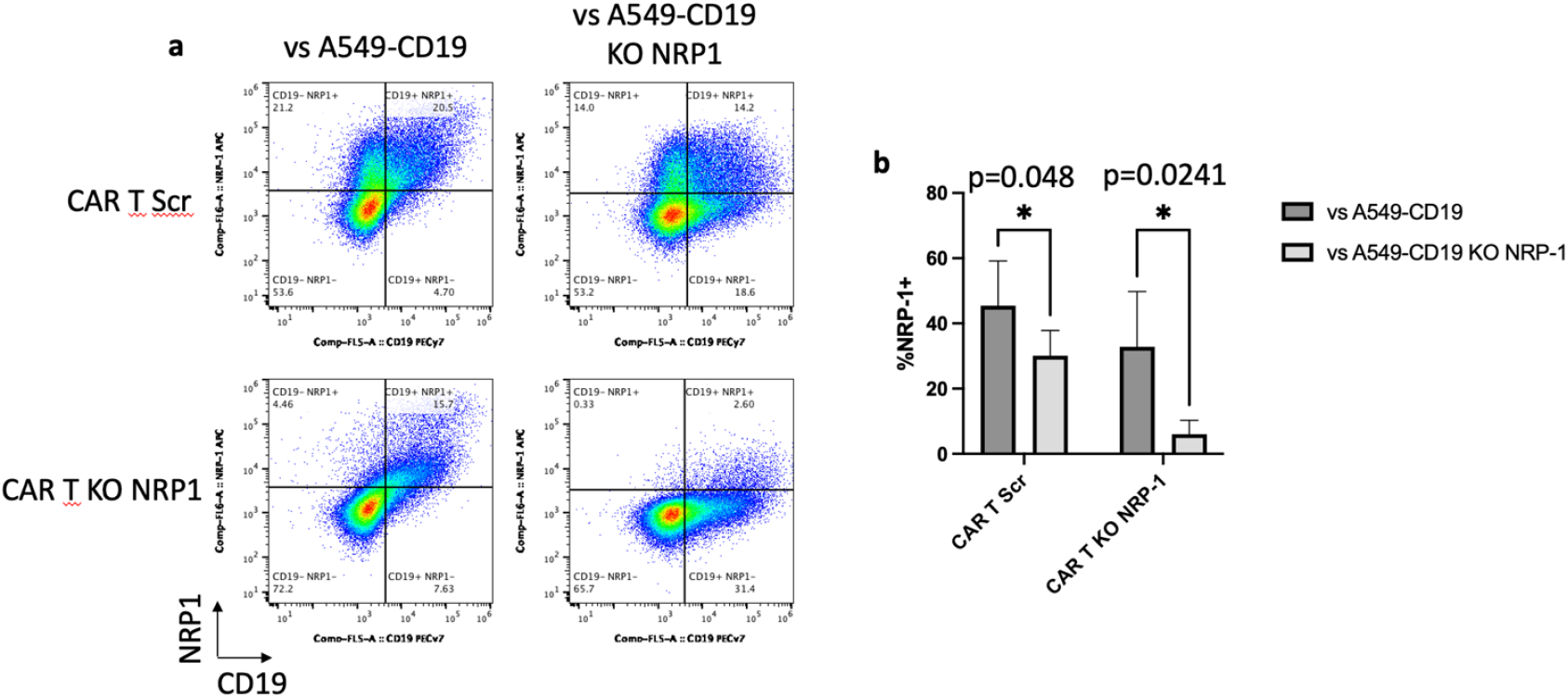
**a.** Representative NRP-1 and CD19 expression on Scr and NRP-1 KO CAR T cells after 24-hour co-culture with A549-CD19 and A549-CD19 NRP-1 KO target cells. **b**. Mean NRP-1 expression on Scr and NRP-1 KO CAR T cells after 24-hour co-culture with A549-CD19 and A549-CD19 NRP-1 KO target cells. Data from 4 independent experiments. Bars indicate standard errors.

This phenomenon also occurred in Scr CAR T cells, which express NRP-1 intrinsically when activated: significantly higher levels of NRP-1 were detected on these effectors after co-culture with NRP-1^+^ target cells than with NRP-1-target cells, without any change in other phenotypical markers or in activation level (**Figure 6a**, top panels, and **6b**).

This phenomenon of transfer and display of plasma membrane proteins from one cell to another is called trogocytosis. Tumor-specific T cells and CAR T cells can acquire tumor antigens by trogocytosis, or cell “nibbling”, upon formation of an immunological or cytotoxic synapse. This can put them in danger of being recognized by other cytotoxic T cells and resulting in fratricide killing within T cell populations; it can also enable tumor escape by stripping antigens from tumor cells (Hamieh *et al*., 2019; Schoutrop *et al*., 2022).

### Tumoral NRP-1 impairs the transfer of target antigen to CAR T cells

As NRP-1 is expressed both by the target cells and the effector cells in this model, we sought a better marker for detecting the transfer of membrane proteins from the former to the latter *via* trogocytosis. CD19 appeared as the most appropriate candidate, being the target of the CAR T cells and thus necessarily involved in the immunological/cytotoxic synapse. CD19 was indeed detectable in a significant proportion of CAR T cells, both Scr and NPR-1 KO, after co-culture with either NRP-1^+^ or NRP-1-A549-CD19 cells (**Figure 6**). While the quantities of transferred CD19 and the resulting proportions of CD19^+^ effector cells varied considerably depending on the T cell donor, the proportion of these CD19^+^ effectors was consistently higher in co-culture with A549-CD19 NRP-1 KO target cells than with NRP-1^+^ target cells (**Figure 7**). This difference was significant in NRP-1 KO CAR T cells, though the same trend was observed in Scr CAR T cells. Thus, more CD19 protein was transferred from target to effector cells in the absence of NRP-1 on the target cells, suggesting that tumoral NRP-1 partially inhibits trogocytosis in this model. An extensively-documented consequence of tumor antigen trogocytosis by various immune cells is the risk of fratricide by other immune cells targeting the same antigen (Li *et al*., 2022). In CAR T cells in particular, strong interactions between a high-affinity CAR and the tumor antigen result in extensive antigen transfer to the effector cell. This in turn provokes a loss in efficacy of these CAR T cells as they are targeted and killed by other CAR T cells. Reducing CAR affinity for their antigen has shown promise for limiting fratricide, which is counter-productive for adoptive immunotherapy (Hamieh *et al*., 2019).

**Figure 7.**
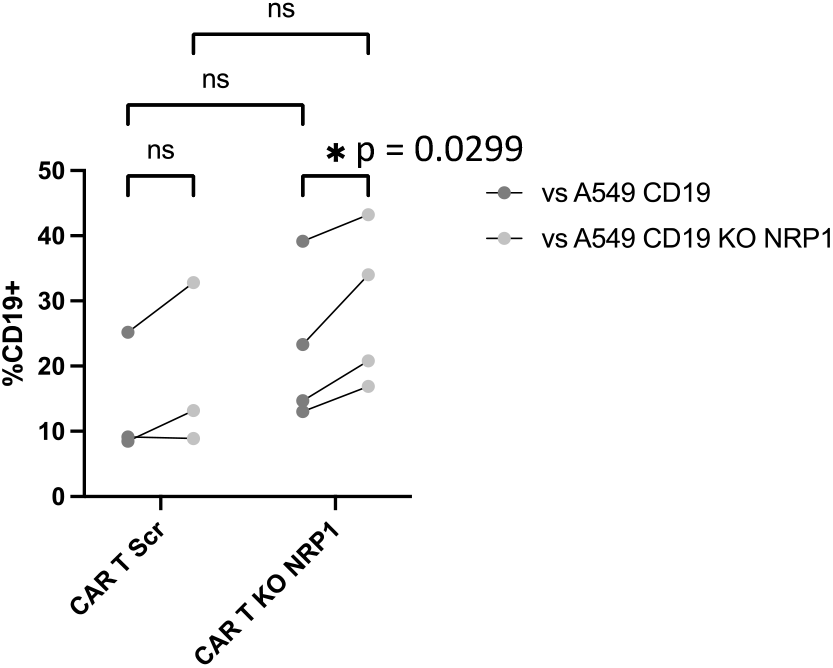
CD19 expression on Scr and NRP-1 KO CAR T cells after 24-hour co-culture with A549-CD19 and A549-CD19 NRP-1 KO target cells. Data pooled from 4 independent experiments.

Studying the numbers of surviving effector cells after co-culture with A549-CD19 targets pointed towards a possible involvement of CAR T cell fratricide. A549-CD19 cells were plated in flat-bottomed wells and allowed to attach for 5 hours, then CAR T cells were added and co-cultured for 24 hours. The CAR T cells were then collected in two steps: first the supernatant only, without disturbing the cells at the bottom of the well (unflushed condition), then the wells were flushed with PBS, collecting the more tightly attached effector cells (flushed condition).

In both conditions, the proportion of CD19^+^ CAR T cells and the amount of CD19 transferred to effector cells during co-culture correlated negatively with the approximate number of live effector cells remaining in culture after 24 hours (**Figure 8**). Thus, higher proportions of CD19^+^ effector cells correlated with lower effector cell survival. We hypothesize that this is due to fratricide death of CD19^+^ CAR T cells. However, we cannot rule out the possibility that CD19^+^ effector cells were more likely to die as a result of stronger activation, with high CD19 expression indicating strong effector function followed by activation-induced cell death (AICD).

**Figure 8.**
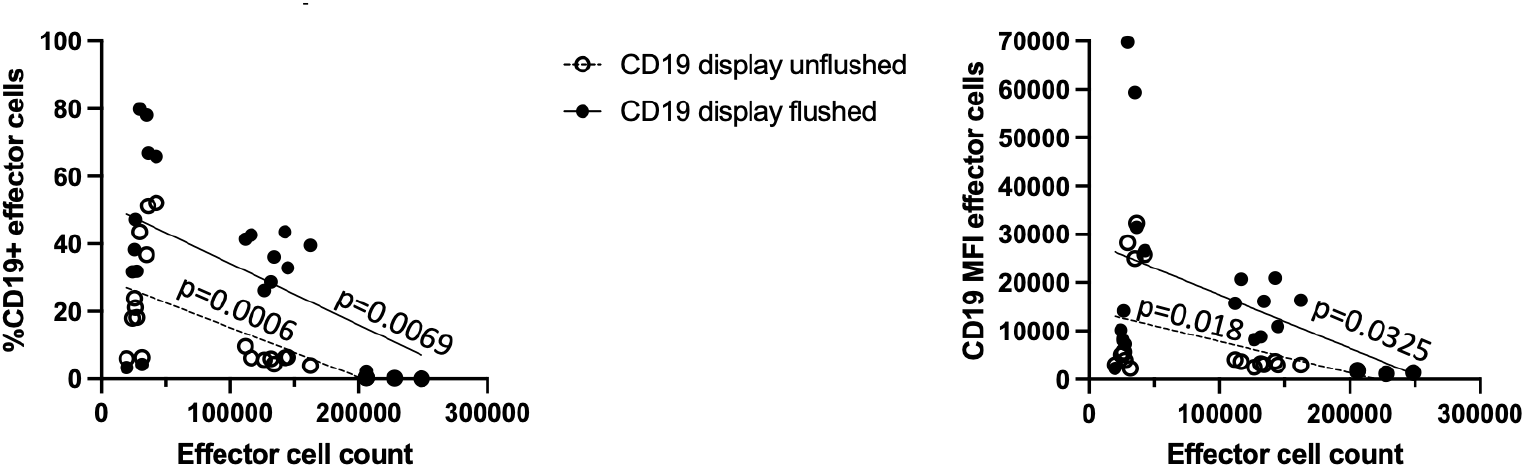
Approximate live T cell count after 24-hour co-culture with A549-CD19 target cells vs CD19 display by these T cells (percentage of total cells displaying CD19 in left panel; MFI or density of CD19 in right panel). P values under 0.05 indicate significantly non-zero slopes, *i*.*e*. a correlation between the two sets of values. Data pooled from 4 independent experiments.

### Antigen density on target cells is not discernably reduced from trogocytosis

To determine whether the CAR T cells were stripping significant amounts of CD19 from target cell plasma membranes, thus potentially facilitating tumor escape by reducing their immunogenicity, we analyzed CD19 expression by the target cells after co-culture with CAR T cells, using flow cytometry.

CD19 expression on target cells was not reduced after co-culture, either when considered as the percentage of CD19-positive cells or as CD19 density at the cell membrane, represented by mean fluorescence intensity (MFI) (**Figure 9**).

**Figure 9.**
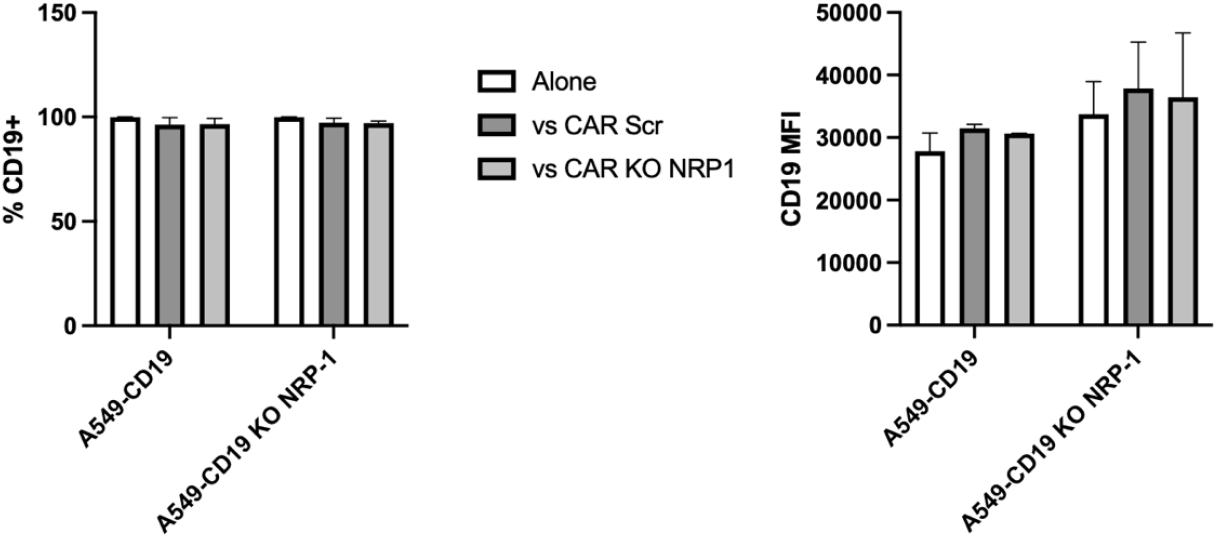
CD19 expression by A549-CD19 and A549-CD19 KO NRP-1 target cells after 24-hour co-culture with Scr or NRP-1 KO CAR T cells (percentage of total cells expressing CD19 in left panel; MFI or density of CD19 in right panel).

This indicates that the target cells were not detectably stripped of antigen, making this an unlikely method of tumor escape in this *in vitro* model.

### NRP-1 has no effect on target cell-CAR T cell conjugate formation

Since a synapse between antigen-presenting and effector cells is required for trogocytosis to occur, we wondered whether target cell-effector cell conjugates would form faster or more abundantly in conditions where we had observed more trogocytosis, and whether this could explain the different target cells’ susceptibility to CAR T cell-mediated killing.

To this end, we co-cultured A549-CD19 NRP-1^+^ or NRP-1^-^ target cells with Scr or NRP-1 KO CAR T cells for 3 hours, to allow abundant time for synapse formation, but not enough for extensive cytotoxicity. The target cells expressed blue fluorescent protein (BFP), while the effectors were either pre-stained with carboxyluorescein succinimidyl ester (CFSE), a fluorescent cell dye, or incubated after co-culture with a fluorescent anti-CD3 antibody, so that the two cell types could be distinguished in flow cytometry. Events that were simultaneously positive for both markers represented conjugates between target and effector cells, and were indeed enriched in multiplets (events corresponding to several cells detected simultaneously by the low cytometer) compared to either single-positive population.

The number of double-positive events was normalized to co-cultures of CAR T cells with wild-type A549 cells, which do not express CD19 and thus constitute a baseline of accidental or transient cell-cell contacts, or rare recognition by a specific TCR, but not by the transduced CAR (**Figure 10**).

**Figure 10.**
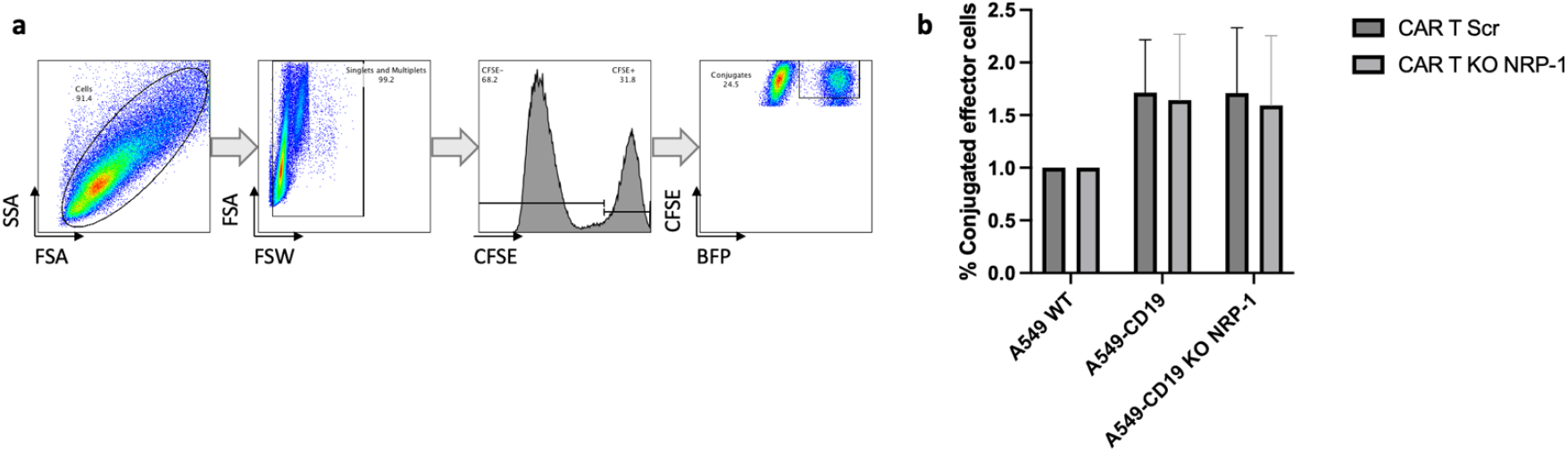
**a.** Flow cytometry gating strategy for identification of effector-target cell conjugates. **b**. Percentage of effector cells conjugated with target cells.

We found no significant differences in numbers of conjugates in the presence or absence of NRP-1 expression, either on the target or on the effector cells (**Figure 10**). It is likely that the influence of NRP-1 on conjugate formation is too small to be detected efficiently using this method.

## Discussion

NRP-1 is a versatile protein that plays multiple roles in the immune response to cancers, but also in the genesis and development of cancers themselves. While it occupies an inhibitory function in conventional T cell responses to tumor cells, we found no such inhibitory role when expressed by CAR T cells, and even some evidence to suggest that the CAR T cell response may be less durable *in vivo* in the absence of NRP-1 expression. This corroborates mounting evidence that CAR T cell biology and function differ more fundamentally that previously thought from that of conventional T cells. NRP-1 has already been shown to be a multifaceted membrane protein with very context-dependent roles, and it is thus perhaps not surprising that it would have different effects in conventional T cells and CAR T cells.

While NRP-1 has been identified in multiple cancers as a pro-angiogenic, tumorigenic, pro-invasive, and poor-prognosis factor, this is the first study to show that its expression specifically protects tumor cells from attack by CAR T cells. CAR T cells directed against A549-CD19 spheroids were less effective against NRP-1^-^ target cells than against NRP-1^+^ target cells, with no discernable differences in intrinsic tumor cell proliferation based on NRP-1 expression.

We speculate that constitutive NRP-1 expression by the target cells used in our model may have dampened the effect of NRP-1 neutralizing antibodies by binding anti-NRP1 themselves. The dose of anti-NRP1 was chosen based on previous work ascertaining total NRP-1 neutralization in T cells at this concentration, but with different target cells (Rossignol *et al*., 2022). Antibody capture by A549 cells may explain their lack of effect on CAR T cell efficacy, hence our choice to rely on cell-specific genetic deletion of NRP-1 in subsequent experiments.

Interestingly, although NRP-1 genetic deletion in target cells made them more vulnerable to CAR T cell cytotoxicity, simultaneous blockade of NRP-1 on the target and effector cells using antibodies had no effect. This suggests that NRP-1 may have contradictory functions that cancel each other out in this system, respectively protecting tumor cells and enhancing CAR T cell activity. Upon indiscriminate NRP-1 blockade, this balance was maintained, but with an excess of CAR T cells at the highest tested dose, the saturation of the antibody may have tipped the balance in favor of non-NRP1-blockaded CAR T cell activity.

NRP-1 and the target antigen CD19 were both transferred to CAR T cells in significant quantities after co-culture with their targets, through a process called trogocytosis. Trogocytosis requires the formation of a synapse between cells, usually involving the TCR, but has more recently been observed in CAR T cells. CAR T cells displaying their own target antigen are at risk of being targeted and killed by fellow CAR T cells through mutual recognition and fratricide. This is a likely explanation for the negative correlation we identified between CD19 acquisition by CAR T cells and their survival after co-culture with target cells.

However, the detection of target cell membrane proteins on T cells is not necessarily a marker for an immunosuppressive phenomenon, as they become targets for fratricide; on the contrary, it can signify effective tumor cell killing, as the acquisition of target cell membrane molecules is a marker that precedes cell-mediated cytotoxicity (Horner *et al*., 2007). This possibility makes it difficult to rule out AICD as a cause for reduced survival among CAR T cells that acquired CD19 after co-culture with target cells. A more definitive way to ascertain whether CD19 acquisition by CAR T cells results in fratricide would be to harvest and isolate these CAR T cells after co-culture with target cells and incubate them alone for 24 to 48 hours. Remaining live CAR T cells could then be counted precisely, along with cell death marker analysis using low cytometry, to determine whether trogocytosis of CD19 leads to CAR T cell fratricide in this model.

The phenomenon of trogocytosis had several potential ramifications to explore: irst, we sought to determine whether this membrane protein transfer was NRP-1-dependent, and if so, whether it was a target cell factor or an effector cell factor. We found that NRP-1 expression by the tumor cell line impeded CD19 transfer to CAR T cells, which seems to indicate that NRP-1’s protective effect on tumor cells does not occur through promoting CAR T cell fratricide. In this case, the acquisition of CD19 by CAR T cells may simply be an indicator of their cytotoxic activity, as described above (Horner *et al*., 2007) – an activity that is dampened by tumoral NRP-1 expression.

Another consequence of trogocytosis is the stripping of these antigens from tumor cell membranes, which can promote their escape from immune surveillance (Hamieh *et al*., 2019; Schoutrop *et al*., 2022). We observed no reduction in CD19 antigen density on the target cell plasma membrane regardless of NRP-1 expression, suggesting that antigenic escape is unlikely to be a mechanism for NRP-1-mediated CAR T cell inefficacy in this model.

However, there are other likely reasons for this apparent lack of antigen-low tumor escape. First, there is a significant difference in size between CAR T cells and A549 target cells, which intrinsically limits the ability of the effector cells to strip large quantities of CD19 from the target cells’ membranes. Second, CD19 production and turnover at the plasma membrane may outpace the CAR T cells’ trogocytic activity, and the timeframe of the experiment may also have allowed the target cells to replenish CD19 expression at the membrane before observation in flow cytometry.

Furthermore, NRP-1 expression did not appear to affect the formation of conjugates between target and effector cells, despite NRP-1 being previously shown to play a central role in immune synapse formation and durability (Sarris et al., 2008; S.-D. Liu *et al*., 2021). Our study is limited in this regard, as our method of conjugate detection relied on low cytometry, which does not distinguish between two cells that happen to be close or touching, and two cells forming a true, stable synapse. Future studies will require direct observation of effector-target doublets and immunological synapse formation using luorescence microscopy, as well as determining optimal conditions for conjugate formation.

While we were not able to identify a specific mechanism for the protective effect of tumor NRP-1 against CAR T cells, we explored several hypotheses and were able to rule out a number of possible mechanisms. One remaining hypothesis is based on NRP-1’s role in cell-cell adhesion. Indeed, we observed that A549-CD19 NRP-1 KO spheroids appeared to have a more diffuse structure than A549-CD19 NRP-1^+^ spheroids. This looser structure may be more permissive to CAR T cell infiltration, giving the effector cells better access to their targets and promoting anti-tumor activity. NRP-1’s protective effect may stem from its ability to promote tight junctions between the tumor cells, impeding CAR T cell penetration and disruption of the spheroid structure. This direct adhesive function has been documented in prostate cancer (Li *et al*., 2016) and in vascular endothelial cell junctions (Roth *et al*., 2016), and future studies should explore the effects of NRP-1 on tumor cell-tumor cell adhesion in addition to tumor cell-effector cell contacts. Scratch wound migration assays, capillary or trans-well migration assays, or any number of cell traction force assays, such as those outlined in Ungai-Salánki et al. (2019), could yield important clues as to the involvement of NRP-1 in tumor cell protection from immune effector attack. NRP-1’s role in tumor cell adhesion and migration could be a key focus point for future efforts to improve CAR T cell therapy in solid tumors.

## Materials and methods

### Primary cells and CAR T cell manufacturing

Peripheral blood mononuclear cells (PBMCs) were isolated from healthy donor apheresis products using a Ficoll density gradient medium and centrifugation. The PBMC ring was collected and washed in RPMI medium, then the cells were counted. T cells were then isolated using the human Pan T Cell Isolation Kit (Miltenyi), according to manufacturer instructions. Through this negative selection method, we obtained T cells untouched by antibodies, which were then frozen in 10% dimethylsulfoxide (DMSO) until needed.

### Activation (D0)

T cells were thawed in pure fetal calf serum (FCS), then washed in their culture medium, X-VIVO (Lonza) supplemented with 5% human serum and 50 µM β- mercaptoethanol. The cells were then incubated at 1 million cells/mL medium for 4 to 5 hours at 37°C/5% CO_2_ before counting and activating using CD3/CD28 Dynabeads (Thermo Fisher). The beads were washed twice with PBS, then added to the cells in a 1:1 ratio and plated together at a concentration of 1 million cells/mL, 500 µL per well in a 24-well plate. Cells were incubated for approximately 24 hours at 37°C/5% CO_2_.

### Viral transduction (D1-D2)

The CD19 CAR construct, containing an anti-CD19 scFv, a 4-1BB costimulatory domain, and the CD3ζ chain, along with an EF1α promoter, was coded in lentiviral vector rLV.EF1.19BBz (Flash Therapeutics). This lentiviral vector was transduced into the stimulated T cells at MOI 10 in 500 µL/well X-VIVO medium with 4µg/mL polybrene, then incubated for 14-16 hours at 37°C/5% CO_2_. The viral supernatant was then carefully removed and replaced with fresh X-VIVO medium containing 100 U/mL IL-2, to a final volume of 2 mL/well.

*CRIPSR-Cas9 nucleofection (D3-D4):* CD3/CD28 Dynabeads were removed magnetically and washed twice to ensure no cells remained attached. T cells were then electroporated with pre-complexed Cas9/gRNA ribonucleoproteins (RNPs), in a molecular ratio of 1:3. 10 µg Cas9 (IDT) were used for 2 million cells, i.e. 61 pmol Cas9 complexed with 183 pmol gRNA per nucleofection. A mix of gRNAs was used, targeting exon 2 of the Nrp-1 gene: CAGGAGAUGUAAGGUACCCG, CUAUAAAAAUUGAAAGCCCC, ACUUUCAUUGCAGAUAAAUG (Synthego). Alternatively, Mock CAR T cells were generated by transfecting Cas9 + buffer instead of gRNAs, and Scr CAR T cells were produced using Cas9 + scrambled gRNAs instead of gRNAs targeting NRP-1. After electroporation using the Lonza 4DNucleofector and the Lonza P3 Primary Cell Kit according to manufacturer specifications, pre-warmed X-VIVO medium with 500 U/mL IL-2 was immediately added and the cells were incubated at room temperature (RT) for 10 minutes. The cells were then gently transferred to a 96-well plate (2 million cells per well in approximately 300 µL) and incubated overnight at 37°C/5% CO_2_. The cells were split the next day and cultured in 24-well plates, 1 million cells per well in 2 mL X-VIVO medium with 500 U/mL IL-2.

The cells were used in experiments at day 6 since thawing and maintained in culture at 0.5-1 million cells/mL in X-VIVO medium with 100 U/mL IL-2.

### Tumor cell lines

Human lung adenocarcinoma cell line A549 was used as a target for CAR T cells. A549 cells already transduced with CD19, BFP, and luciferase were kindly provided by Sebastian Amigorena’s team at Institut Curie. These cells, along with A549 cells not expressing CD19, were additionally transduced with GFP to be used in real-time imaging of cell killing assays.

NRP-1 knock-out was performed in these cell lines with the same gRNA mix as described previously, using the Lonza SF Cell Line nucleofection kit and 4D-nucleofector. The cells were not sorted, as over 95% reduction in NRP-1 expression was achieved using this method.

All A549 cells were routinely cultured in RPMI medium supplemented with 10% FCS and 1% penicillin-streptomycin, in T75 flasks, and detached with 0.05% trypsin for passaging twice per week.

### Spheroid assays with real-time imaging

1000 A549 cells per well were seeded in ultra-low adhesion (ULA) 96-well plates, in 100 µL RPMI medium, and centrifugated at 150xg for 5 minutes, with low acceleration and brake. Sterile PBS was added in the spaces between the wells to limit evaporation, and the plates were incubated at 37°C/5% CO_2_ for 3 days.

CAR T cells were collected and analyzed by flow cytometry at day 6 since their irst activation, then added directly to the spheroids in doses varying from 200 to 10000 CAR^+^ cells per well, in 100 µL X-VIVO.

The wells were imaged every 4 hours for approximately 1 week in an IncuCyte S3 Live Cell Analysis Instrument (Sartorius).

For further analysis of the cells’ phenotypes after these assays, the medium in each well was gently pipetted up and down to collect cells floating in the supernatant, making sure not to aspirate the solid spheroid, when still present. The spheroids were then washed in PBS then dissociated using 0.05% trypsin, which was saturated with an excess of FCS before analysis by flow cytometry.

### Trogocytosis assays

1 million A549 cells per well were plated in 6-well plates and incubated for 4-5 hours at 37°C/5% CO_2_ to adhere, then the same number of CAR T cells was added and the plates were incubated for 24 hours at 37°C/5% CO_2_.

The cells were collected and analysed the next day. Similarly to collection of cells in spheroid assays, the supernatant was gently pipetted to collect cells in suspension or weakly attached, then the attached cells were washed with PBS and detached with trypsin. The two cell populations were stained and analysed separately by flow cytometry.

### Conjugate assays

CAR T cells and A549 target cells were incubated together in a 1:1 ratio in 96-well round-bottom plates for 3 hours at 37°C/5% CO_2_. To avoid disturbing any attached cells, the plate is directly washed with PBS 2% FCS and centrifugated, and resuspension in low cytometry antibody mixes is done with minimal agitation. Each well is slowly pipetted precisely the same number of times to resuspend the cells, to avoid introducing any accidental disparities in conjugate numbers between conditions.

### Flow cytometry

After collection, cells were resuspended in cold PBS with 2% FCS. They were stained for 20 minutes at 4°C using combinations of the following antibodies and their corresponding control isotypes (all from Biolegend, unless otherwise specified). For CD19 CAR staining, a primary antibody conjugated to biotin was added first, incubated for 20 minutes at 4°C, then washed, then the secondary antibody targeting biotin was included in the rest of the antibody mix.

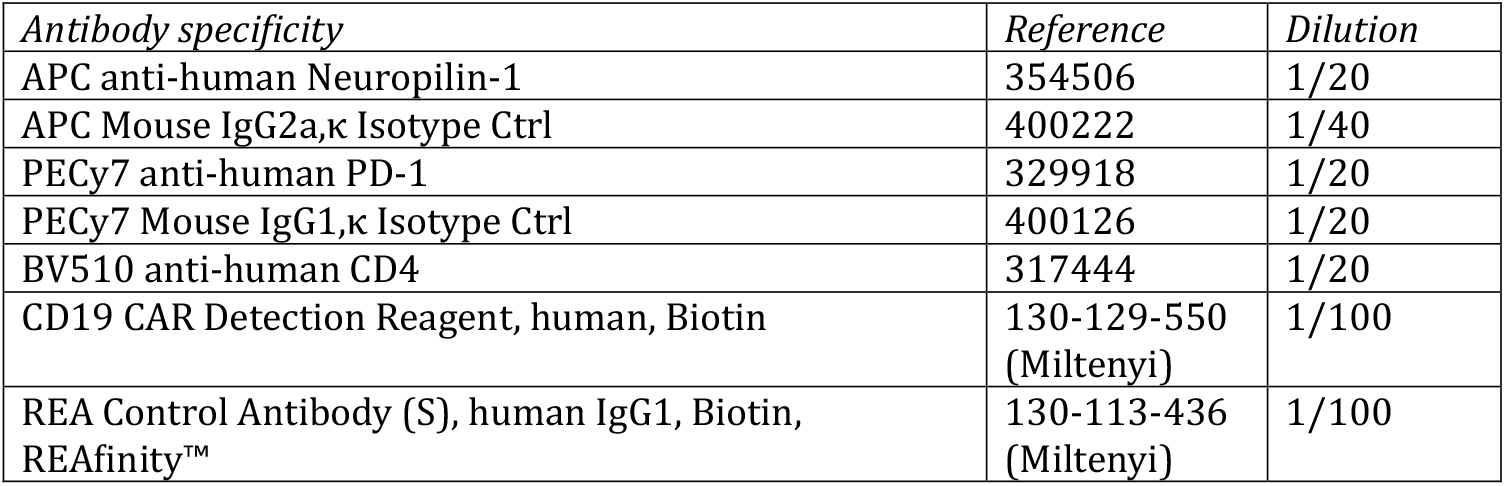
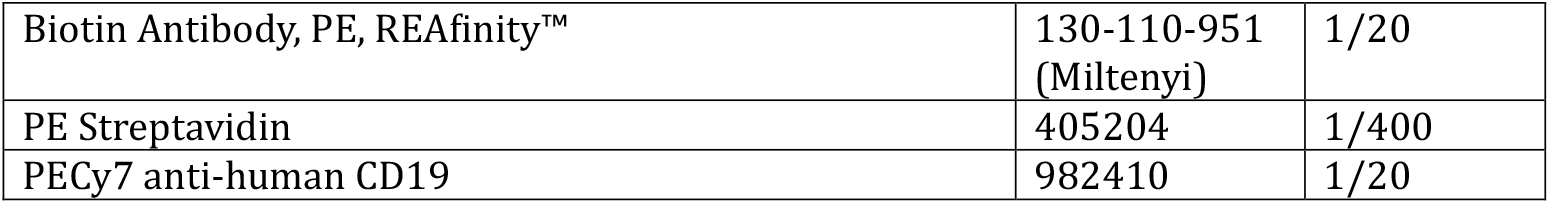

| *Antibody specificity* | *Reference* | *Dilution* |
| --- | --- | --- |
| APC anti-human Neuropilin-1 | 354506 | 1/20 |
| APC Mouse IgG2a,κ Isotype Ctrl | 400222 | 1/40 |
| PECy7 anti-human PD-1 | 329918 | 1/20 |
| PECy7 Mouse IgG1,κ Isotype Ctrl | 400126 | 1/20 |
| BV510 anti-human CD4 | 317444 | 1/20 |
| CD19 CAR Detection Reagent, human, Biotin | 130-129-550<br>(Miltenyi) | 1/100 |
| REA Control Antibody (S), human IgG1, Biotin, REAfinity™ | 130-113-436<br>(Miltenyi) | 1/100 |
| Biotin Antibody, PE, REAfinity™ | 130-110-951<br>(Miltenyi) | 1/20 |
| PE Streptavidin | 405204 | 1/400 |
| PECy7 anti-human CD19 | 982410 | 1/20 |

### *In vivo* assays

Young adult NSG mice were injected intravenously with A549-CD19 cells expressing luciferase, which were then left to home to the lungs and form tumors for 3 weeks. After 3 weeks, day 6 Mock or NRP-1 KO CAR T cells were injected intravenously. Tumor size was monitored weekly by intraperitoneal administration of luciferin (150 µg per gram of body weight) and luminescence imaging using an IVIS imaging system.

